# scRecover: Discriminating true and false zeros in single-cell RNA-seq data for imputation

**DOI:** 10.1101/665323

**Authors:** Zhun Miao, Jiaqi Li, Xuegong Zhang

## Abstract

High-throughput single-cell RNA-seq (scRNA-seq) data contains excess zero values, including those of genes not expressed in the cell, and those produced due to dropout events. Existing imputation methods do not distinguish these two types of zeros. We present a modest imputation method scRecover to only impute the dropout zeros. It estimates the zero dropout probability of each gene in each cell, and predicts the number of truly expressed genes in the cell. scRecover is combined with other imputation methods like scImpute, SAVER and MAGIC to fulfil the imputation. Down-sampling experiments show that it recovers dropout zeros with higher accuracy and avoids over-imputing true zero values. Experiments on real data illustrate scRecover improves downstream analysis and visualization.

## 1 Introduction

Single-cell RNA sequencing (scRNA-seq) is becoming an essential technology in current genomics studies especially in the Human Cell Atlas project^1^. scRNA-seq data usually contain more zeros comparing to bulk RNA-seq data^2–4^. The excessive zeros can come from true zero expressions^5,6^ or technique noises caused by factors such as low mRNA capturing rate^7,8^. The latter is often called “dropout” zeros^9^. Too many dropout zeros may cause problems for downstream analysis since they may distort the structure and characteristics of gene expression profiles.

Several methods have been developed to impute the zero values in scRNA-seq matrixes based on information from other genes and other cells^10^. For example, scImpute^11^ imputes zeros by borrowing information from similar cells using gene sets with lower dropout probability. SAVER^12^ uses gene-gene relations to estimate expressions across cells with a Bayesian approach. MAGIC^13^ calculates cell-cell affinities and imputes gene expressions using exponential Markov matrix. scRMD^14^ tries to recover true expression values through robust matrix decomposition and ALRA^15^ solves this problem with low-rank approximation. netNMF-sc^16^ combines gene interaction network and non-negative matrix factorization (NMF) to reconstruct count matrix and improve downstream clustering. Incorporating statistical models into deep-learning framework, DCA^17^ presents a deep count auto-encoder to impute missing values.

Most of those methods impute all the zero values and also alter the non-zero values. As gene transcription is an on-off statistical process in each single cell^5^, many of the zeros can be true zeros of genes that are not transcribed in the cell. Imputing all these zeros to some non-zero values can introduce severe noise or artifacts. This can be seen as “over-imputation”. The methods scImpute and ALRA do not force all zeros to some non-zero values. But they also do not provide the feature of distinguishing true zeros from dropout zeros. scImpute determines a set of genes across all cells that should be imputed according to manually set threshold on dropout probabilities, and ALRA sketches a rough range of impute-free genes based on the symmetry assumption that negative values after low-rank approximation correspond to true zeros. It is desirable to have a modest imputation method that can distinguish true zeros from dropout zeros in the scRNA-seq expression matrix, and do imputation only on those dropout zeros.

In our work on modeling scRNA-seq data with zero-inflated negative binomial (ZINB) distribution, we found that it is possible to estimate the probability of a gene with zero expression in a cell to be a true zero or dropout zero. On the other hand, using the model of species accumulation curve^18^, we can also predict the number of true zeros in a single cell if there were no dropout events. Based on these theoretical work, we proposed a modest imputation method and developed an R package scRecover to recover dropout zeros in scRNA-seq data while keeping the true zeros unchanged. scRecover is applied on top of existing imputation methods such as scImpute, SAVER or MAGIC to implement modest imputation. We conducted a series of experiments using down-sampling scRNA-seq data of relatively high coverage, as well as real scRNA-seq data. Results showed that scRecover could accurately locate dropout zeros and improve the separation of cell clusters in the downstream analysis.

## 2 Methods

### 2.1 Estimating dropout probabilities of genes

ZINB distribution is a mixture of negative binomial (NB) distribution and constant zeros^19^. The probability mass function (*pmf*) of ZINB distribution is:

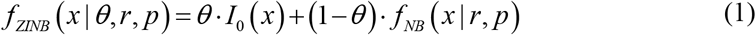

where θ is the proportion of constant zeros of gene g in each cells, *I*_0_(*x*) is an indicator function which equals to 1 for *x* = 0 and 0 for other values, *f*_*NB*_ is the *pmf* of the NB distribution, r is the size parameter and p is the probability parameter of the NB distribution. The zeros in expression matrix consist of constant zeros (true zeros) and zeros from NB part (dropout zeros)^20^.

We chose ZINB distribution to depict the read counts and dropout zeros. For one gene, its expression across cells before mRNA capturing is modeled as:

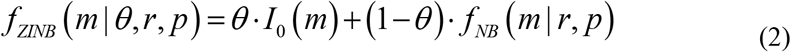

After mRNA capturing,

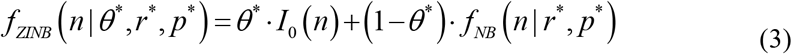

Referring to the demonstration in DEsingle^20^, the relations between parameters are:

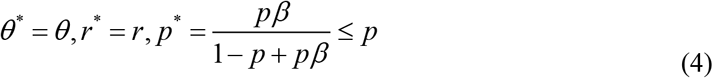

where β is the random capture efficiency of mRNA capture procedure.

The ZINB distributions with different random capture efficiencies are shown in <u>**Supplementary Figure 1**</u>.

Considering cell types, for each gene i, it’s expression in cell subpopulation k is modeled as a random variable 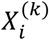 with density function:

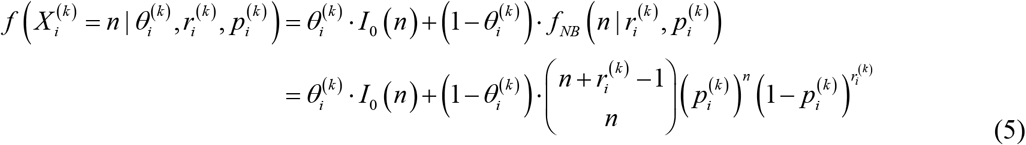

Then we calculate the dropout probability of zeros in gene i, which belongs to subpopulation k:

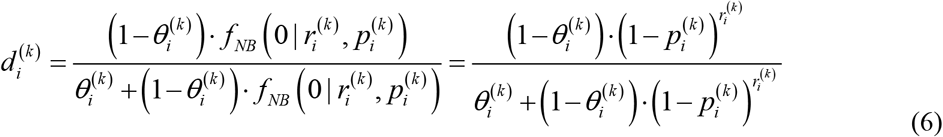

### 2.2 Estimating expressed gene number in each cell

Next, we try to estimate dropout zero number in each cell. We summarized this task as predicting the total category number using unsaturated random sampling, which samples partial population of some categories.

We borrowed the idea of the species accumulation curves and the Good-Toulmin series^18^, which are used to estimate the total number of species in a certain area. The Good-Toulmin series is the unbiased nonparametric Bayesian estimation:

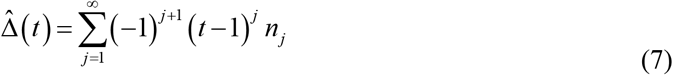

where t is the relative sampling depth, *n*_*j*_ is observed number of species j.

To deal with the sparsity of Good-Toulmin series when t > 2, Daley and Smith proposed rational function approximation^21^ to ensure the accuracy and stability:

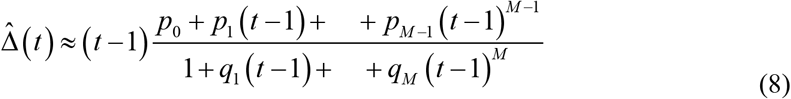

In 2015, Deng and Smith improved the rational function approximation through continued fraction and developed an R package preseqR^22^. Since estimating expressed gene number from expression profile containing dropout zeros is quite similar to estimating species categories by unsaturated sampling from the total population, we employed preseqR to calculate the expressed gene number then obtained dropout zero number X in each cell.

To evaluate the performance of preseqR on scRNA-seq data, we used mouse retina cells data^23^ and down-sampled a proportion of reads in each cell from the original data. Then we performed preseqR and compared the predicted expressed gene number with that of original data. By changing down-sampling rate, we obtained two curves for the predicted number and true number of expressed genes. We randomly chose results of 4 cells, which are shown in <u>**Supplementary Figure 2**</u>. Since two curves are very close with a large overlap, preseqR could make accurate predictions for expressed gene number.

With the dropout probability of each zero value and the dropout zero number X in each cell, we could rank the dropout probabilities in each cell in descending order and identify the top X zeros as dropout zeros in the statistical sense.

### 2.3 Imputing dropout zeros

scRecover employs 3 popular imputation methods: scImpute, SAVER and MAGIC, but it is independent from them. We could view it as a 2-leayers imputation framework. First, we run these intermediate imputation methods on the observed matrix. After each run, scRecover outputs 3 different imputed matrixes based on the results of scImpute, SAVER and MAGIC respectively. The workflow of scRecover is shown in Fig.1.

**Fig. 1.**
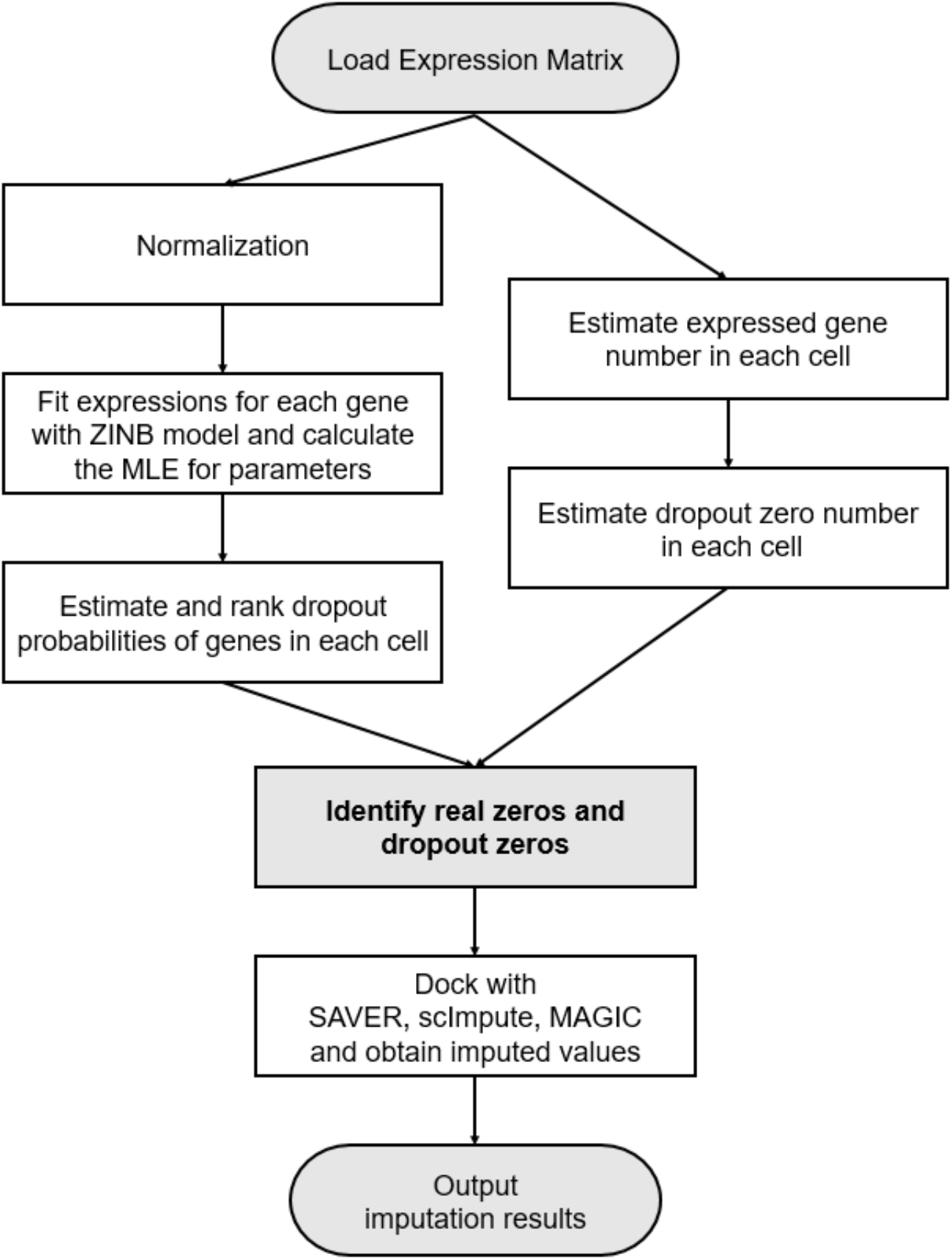
Workflow of scRecover. After loading the expression matrix, scRecover first estimates dropout probabilities of genes and dropout zero number in each cell. Then it could identify true zeros and dropout zeros. scRecover docks with 3 intermediate imputation methods SAVER, scImpute, MAGIC respectively and alters dropout zeros with imputed values, which generates 3 imputed matrixes as the output.

## 3 Results

### 3.1 Performance evaluation on down-sampling data

We compared the prediction accuracy for dropout zero number of scRecover together with scImpute, SAVER and MAGIC on down-sampling data. With different down-sampling rates, we generated datasets from a public scRNA-seq dataset of human preimplantation embryonic cells^24^. For each cell, we removed certain proportion of reads across genes, which is similar to real dropout scenarios. Since the original dataset is generated using SMART-seq protocol with deep sequencing depth, we treated it as reference and evaluate scRecover by comparing recovered matrix with the original one. Following the same procedure, we can also make comparison between scRecover and 3 intermediate imputation methods.

We performed scImpute, SAVER, MAGIC together with scRecover then calculated the accuracy (Fig.2 (a)) and dropout zero number (Fig.2 (b)) on the result of down-sampling datasets with different down-sampling rates. The accuracy is the ratio of correctly-predicted zero number (including true zeros and dropout zeros) over the total zero number in reference dataset. Prediction depth is the ratio of the percentage of reads for prediction over the percentage of reads used, which is a parameter of preseqR package^22^.

**Fig. 2.**
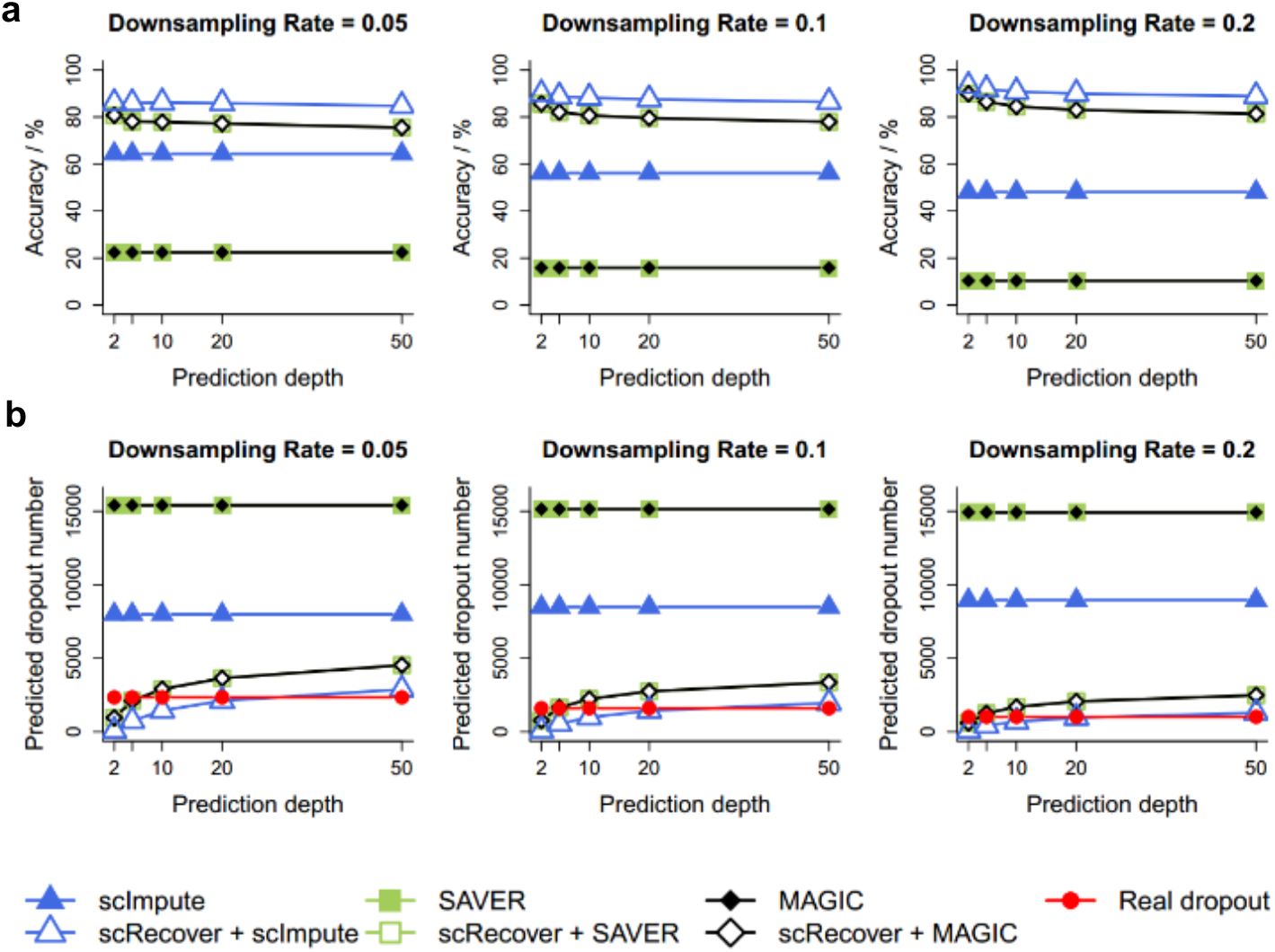
scRecover achieves better performance for true and dropout zero predictions. This plot was generated under parameter Kcluster = 1. **(a)** The relationship between zero prediction accuracy and prediction depth with different down-sampling rates. **(b)** The relationship between dropout zero number prediction and prediction depth with different down-sampling rates.

Zero-prediction accuracy of scRecover plus intermediate imputation methods improves at least 20% compared with those without scRecover, especially for SAVER and MAGIC. As the down-sampling rate increases, the improvement of accuracy becomes more significant. For dropout zero number prediction, results of scRecover plus intermediate imputation methods are much closer to the reference results compared with those without scRecover, even with the increase of prediction depth. Results with other numbers of Kcluster are shown in <u>**Supplementary Figure 3**</u>. On down-sampling datasets, scRecover could not only provide accurate predictions for dropout zeros but also locate them precisely.

### 3.2 scRecover improves clustering analysis

We also assessed the performance of scRecover with clustering analysis on two real datasets. One is 1k heart cells from an E18 mouse downloaded from 10x genomics. The other is human pluripotent stem cells dataset^25^ using SMART-seq protocol. In the 1k mouse heart cells dataset, there is no label for individual cells. Here we treated the cell groups provided by 10x genomics as labels. To evaluate imputation methods, we employed the commonly used analyzing pipeline Seurat^26^ to complete clustering on original data, intermediate imputation results as well as scRecover outputs. If the clustering result using expression matrix after imputation could separate cell groups better, the corresponding imputation method achieves better performance. Differently, the human pluripotent stem cells dataset contains cell labels marked by fluorescence-activated cell sorting (FACS). Better imputation method could group cells with the same label together in clustering analysis.

For both datasets, we drew t-SNE plots for intermediate results of scImpute, SAVER, MAGIC and the outputs after scRecover processing respectively. The plots for two datasets are shown in Fig.3. We also calculated adjusted rand index (ARI) and Jaccard index for each clustering result.

**Fig. 3.**
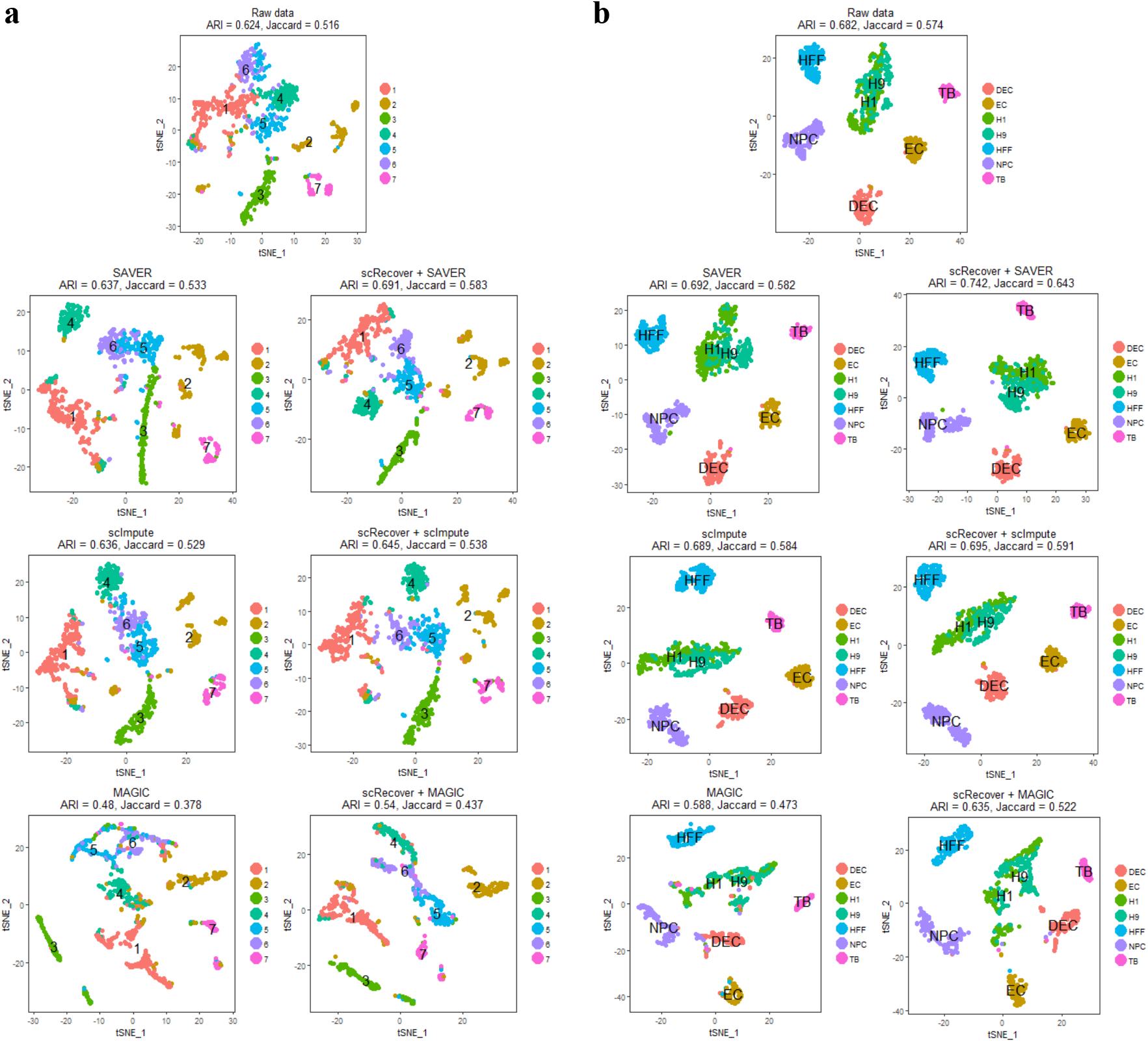
t-SNE plots of mouse heart cells dataset (a) and human pluripotent stem cells dataset (b). For both (a) and (b), on the top is the t-SNE plot for raw data. The left column contains results of intermediate imputation methods including SAVER, scImpute and MAGIC. The right column are results of scRecover on each intermediate imputation method. In (b) we label clusters with the cell type which has the maximum number of cells in this cluster.

For results in Fig.3(a), cluster 5 and 6 are mixed in raw data. Intermediate imputation methods could separate these two clusters to some extent (left column in Fig.3(a)). After scRecover, cluster 5 and 6 are well-separated with increases in both ARI and Jaccard index (right column in Fig.3(a)).

In the analysis provided by 10x genomics (**Data Availability**), CD4 is a marker gene of both cluster 5 and 6. Since nearly all the marker genes of cluster 5 belong to those of cluster 6, these two clusters are much similar to each other. To reveal the slight difference between them, we performed differential expression (DE) analysis only with cluster 5 and 6 samples using DEsingle^20^ and found 3481 DE genes with adjusted *p-value* less than 0.05. We used the top 200 DE genes to do gene ontology (GO) analysis. Results indicated that those DE genes are related to DNA repairing and are highly expressed in cluster 6. By searching studies relevant to DNA repairing in CD4 cells^27–29^, we noticed that high level of protein p27 expressed by gene Cdkn1b in CD4 cells would damage DNA structure and inhibit cell cycles. Also, the expression level of gene Cdkn1b was significantly higher in cluster 6 (adjusted *p-value* 1.87×10^−10^). So these two clusters of CD4 cells are different and scRecover could separate them well.

For results in Fig.3(b), we assign labels for each cluster with the cell type which has the maximum number of cells in this cluster. In Fig.3(b), cells marked as H1 and H9 are mixed in raw data. Intermediate imputation methods could not separate these two clusters well (left column in Fig. 3(b)). After scRecover, H1 and H9 cells become more separated with increasing ARI and Jaccard index (right column in Fig. 3(b)).

H1 and H9 are two different cell lines originated from human embryos^30^. The karyotype is XY for H1 cells while XX for H9 cells. In this human pluripotent stem cells dataset, part of H1 cells were induced to differentiate to DEC, EC, NPC and TB while some H1 cells (n=212), H9 cells (n=162) and human foreskin fibroblasts (HFF, n=159) were used for control.

To further explore the difference between H1 and H9, we again performed gene ontology (GO) analysis using top 200 DE genes found by DEsingle (H1 vs H9, 2651 DE genes in total with adjusted *p-value* less than 0.05). Results indicated that DE genes were enriched in structural constituent of ribosome and rRNA binding. Related study^31^ shows that ribosome protein gene RPS4X is differentially expressed between H1 and H9 and RPS4X is exactly a DE gene found by our study (adjusted *p-value* 3.04×10^−14^). So H1 and H9 human embryonic stem cells are slightly different due to the gender difference, and scRecover improves the separation between these two types of cells.

For both datasets, we calculated average number of expressed, imputed and zero-valued genes across cells for each result, shown in Fig.4. SAVER and MAGIC impute all genes while scImpute leaves part of genes unchanged. After scRecover processing, the number of expressed genes decreases and this is the reason why scRecover makes “modest” imputation. Since there are true zero expressions in scRNA-seq data^5^ and scRecover could locate dropout zeros more accurately, the results of scRecover are more in line with biological knowledge.

**Fig. 4.**
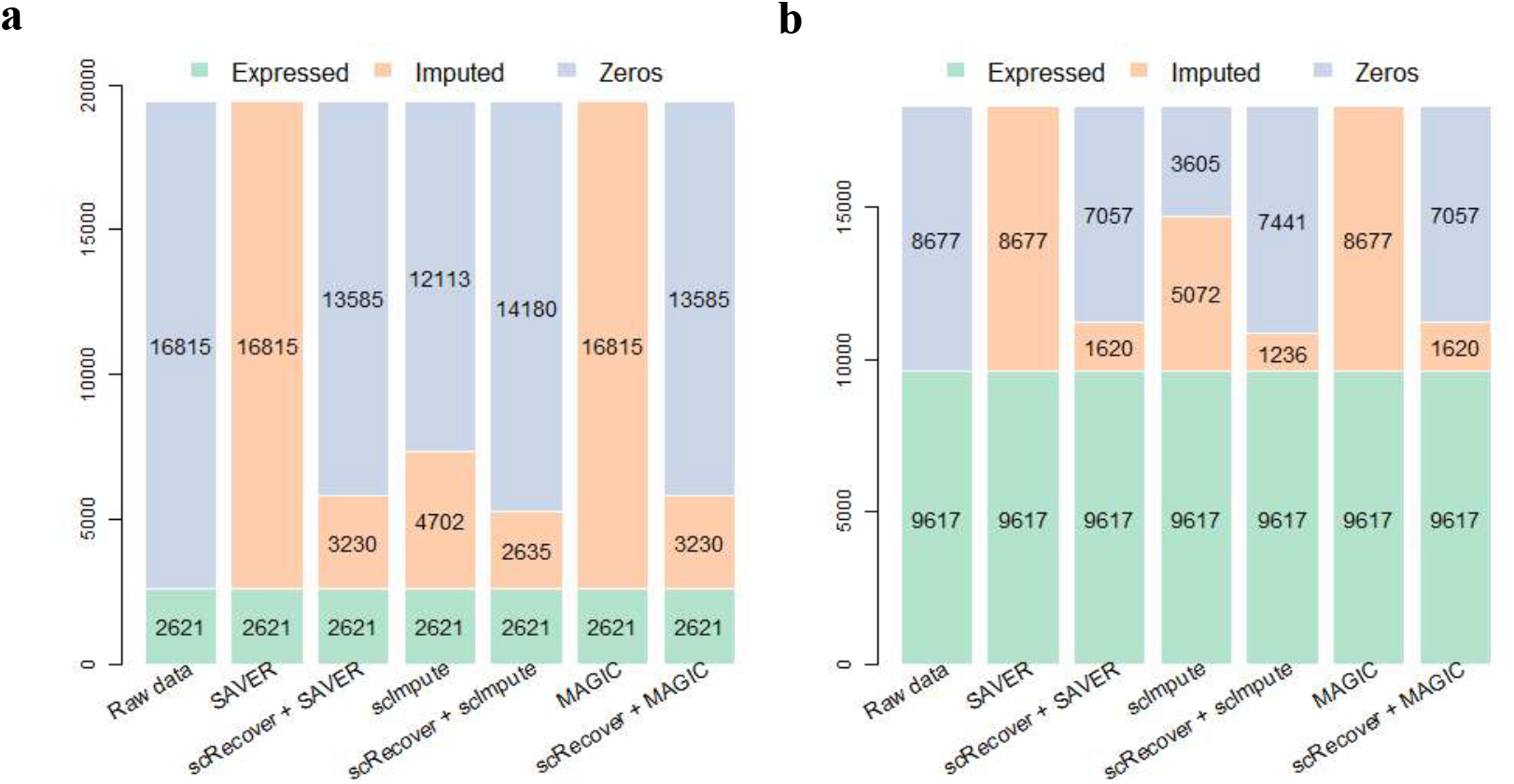
Stacked bar plots for average number of expressed, imputed and zero-valued genes. ScRecover reduces the number of imputed genes and makes “modest” imputation. **(a)** Mouse heart cells dataset. **(b)** Human pluripotent stem cells dataset.

To further analyze the similarity between clustering results and real cell types, we calculated the number of cells that mapped on their cell types for each clustering result in Fig.3(b). We used confusion matrix for visualization (shown in **Supplementary Figure 4**). The results indicate that intermediate imputation methods would cause more mismatches and clustering results of scRecover are closer to ground truth. After scRecover, mismatches reduce and to some extent the characteristics of data are recovered.

## 4 Conclusion

We proposed a modest imputation method and developed an R package scRecover to recover drop-out values in scRNA-seq data. By modeling gene expression in single cells using ZINB model and estimating expressed gene number with species accumulative curve, scRecover could distinguish true zeros from dropout zeros accurately and impute dropout zeros while keeping the true zeros unchanged. The clustering performance improves after scRecover processing compared with those only using intermediate imputation methods such as scImpute, SAVER and MAGIC. We expect that scRecover could provide a new angle for recovering single cell dropout events and make more contributions on downstream analyses.

## Acknowledgement

This work was supported by the National Natural Science Foundation of China (grant numbers 61721003 and 61673231), National Key R&D Program of China grant 2018YFC0910400, Tsinghua-Fuzhou Institute for Data Technology (No.TFIDT2018006) and the CZI HCA pilot project (2017-174037). Conflict of Interest: none declared.

## Supplementary Information

Supplementary figures are available at bioRxiv online.

## Availability and Implementation

The R package scRecover is freely available at Bioconductor and Github https://bioconductor.org/packages/scRecover https://github.com/XuegongLab/scRecover

## Data Availability

All 4 datasets used in this paper are publicly available.

The mouse retina cells datasets could be found at GSE80232.

The human preimplantation embryonic cells datasets could be found at ArrayExpress with accession number E-MTAB-3929.

The E18 mouse 1k heart cells datasets could be found at https://support.10xgenomics.com/single-cell-gene-expression/datasets/3.0.0/heart_1k_v3.

The human pluripotent stem cells dataset could be found at GSE75748.

