## Supplemental Figure 1-4 for "scRecover: Discriminating true and false zeros in single-cell RNA-seq data for imputation"

### Supplementary Figures

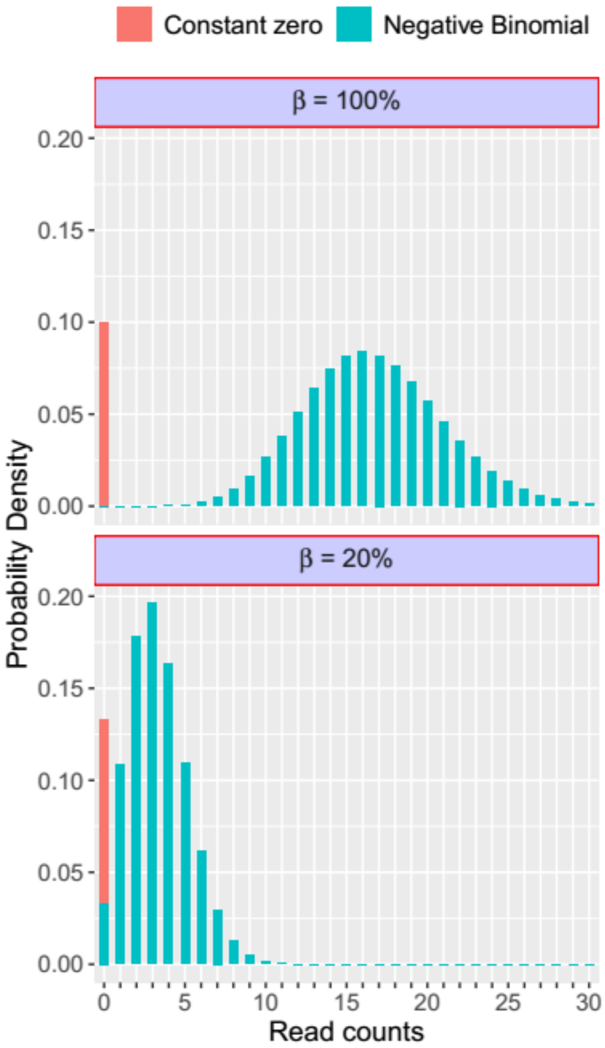


**Figure S1. ZINB distributions with different mRNA random capture efficiencies (β).**

The red bar refers to constant zeros, which are true zeros in scRNA-seq data. The cyan bar refers to negative binomial (NB) distribution. For $\beta = 100\%$, which means all mRNAs have been successfully captured, the distribution is an ideal ZINB. In statistical sense all zeros belong to true zeros. When $\beta$ decreases to 20%, which means 80% of mRNAs are lost, a number of zeros appear in NB distribution. The zeros consist of two parts and those in NB population are dropout zeros.

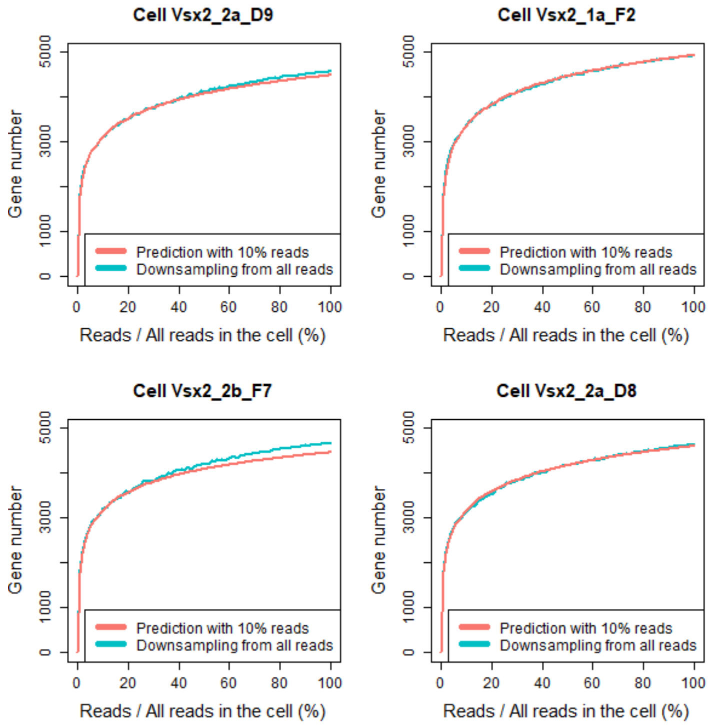


**Figure S2. Evaluation of expressed gene number prediction method.**

We used an R package *preseqR* (Deng et al., 2015) to predict the expressed gene number in each cell. To validate its accuracy, we down-sampled with different proportions of reads and treated the original data as reference. Then we compared the prediction results of *preseqR* (red curve) with the ground truth (cyan curve). The results show that the predicted expressed gene numbers are close to the true number and the accuracy of *preseqR* is validated.

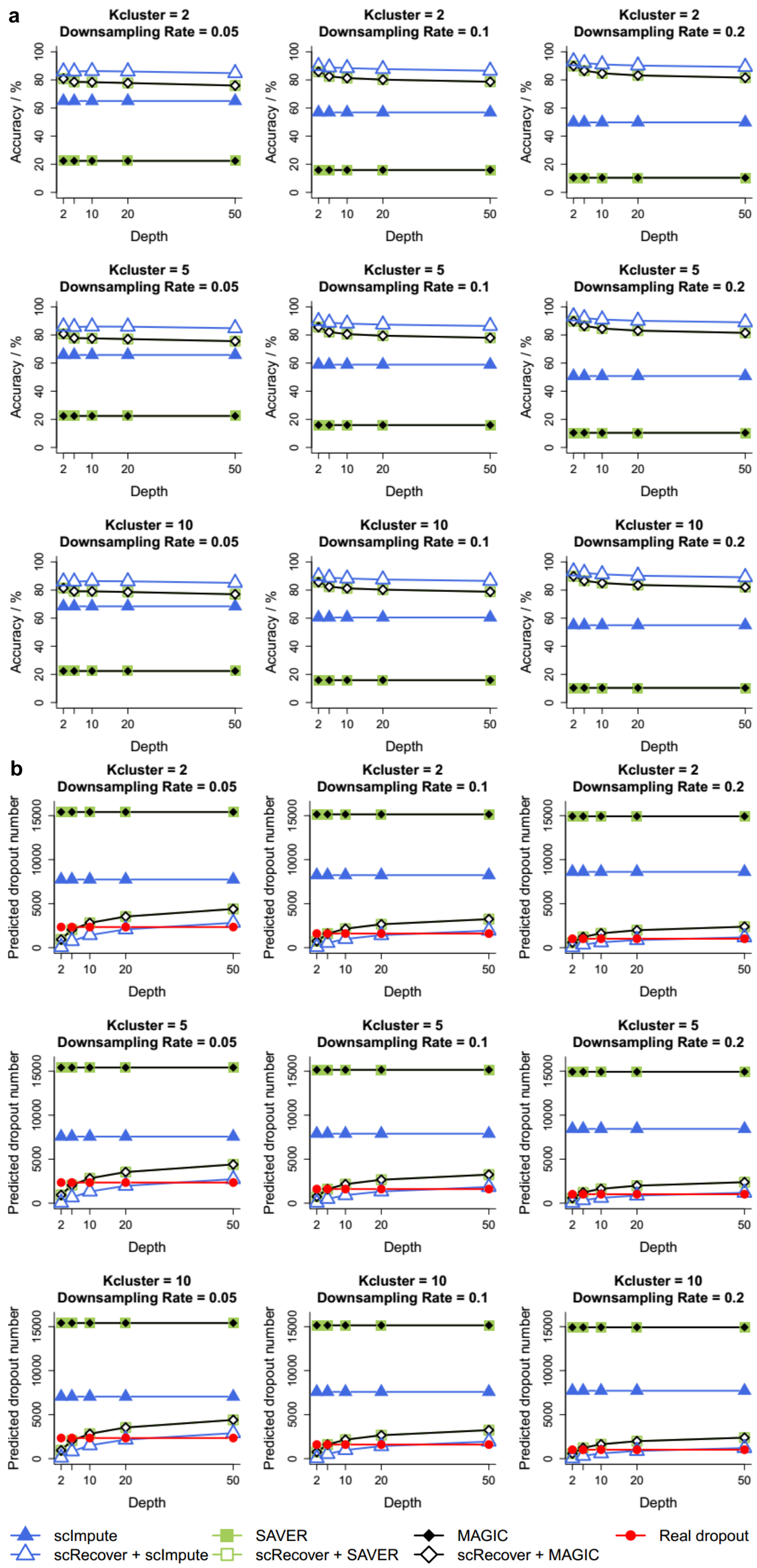


**Figure S3. Zero-prediction accuracy and dropout zero number predictions with different *Kcluster* parameters.**

*Kcluster* is the parameter of estimated cluster number, which is used for pre-clustering. Fig.2 shows the results when *Kcluster* = 1. When it increases to 2,5 and 10 respectively, results are similar.

**(a)** Zero-prediction accuracy of scRecover plus intermediate imputation methods improves nearly or over 20% compared with those without scRecover, especially for SAVER and MAGIC. As the down-sampling rate increases, the improvements of accuracy are more significant.

**(b)** For dropout zero number prediction, results of scRecover plus intermediate imputation methods are much closer to the reference results compared with those without scRecover. scRecover predictions are still closer to reference with the increase of prediction depth.

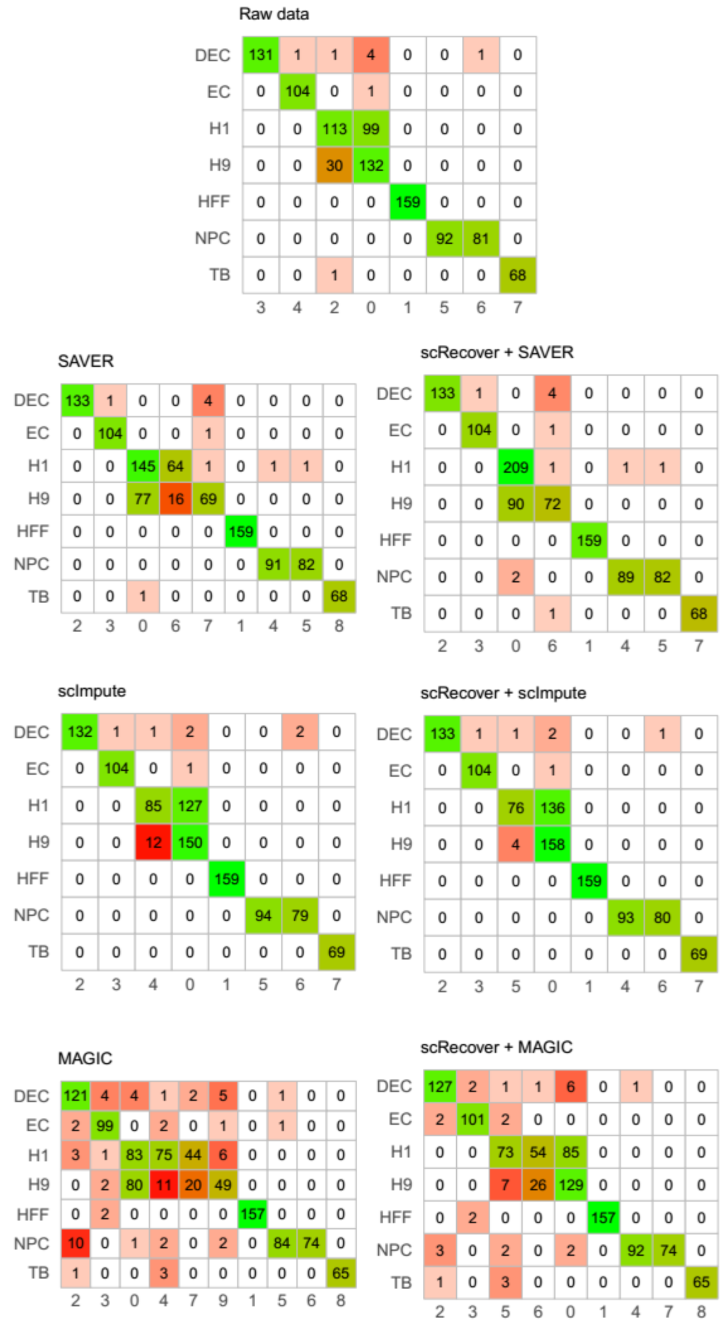


**Figure S4. Similarity analysis between clustering results and real cell types with confusion matrixes.**

We calculated the number of cells that mapped on their cell types for each clustering result in Fig.3(b). The rows of one matrix are the labels assigned for clusters and the columns are real cell types. The number along the diagnose of confusion matrix would be larger if more cells mapped onto their real cell types. The left column contains results of intermediate imputation methods including SAVER, scImpute and MAGIC. The right column are results of scRecover on each imputation method. Colors in each block correspond to the values: white for zero, red for small values and green for large values. Mismatches are mainly between H1 and H9 cells in raw data. Intermediate imputation methods would cause more mismatches (left column), maybe because they could not distinguish true zeros from dropout zeros properly. Altering non-zero values may be another reason for the mismatches in SAVER and MAGIC results. After scRecover, mismatches reduce and to some extent we recover the characteristics of data (right column). The clustering results are closer to ground truth.
